# Whole genome view of the consequences of a population bottleneck using 2926 genome sequences from Finland and United Kingdom

**DOI:** 10.1101/063388

**Authors:** Himanshu Chheda, Priit Palta, Matti Pirinen, Shane McCarthy, Klaudia Walter, Seppo Koskinen, Veikko Salomaa, Mark Daly, Richard Durbin, Aarno Palotie, Tero Aittokallio, Samuli Ripatti, Sequencing Initiative Suomi (SISu) Project

## Abstract

Isolated populations with enrichment of variants due to recent population bottlenecks provide a powerful resource for identifying disease-associated genetic variants and genes. As a model of an isolate population, we sequenced the genomes of 1463 Finnish individuals as part of the Sequencing Initiative Suomi (SISu) Project. We compared the genomic profiles of the 1463 Finns to a sample of 1463 British individuals that were sequenced in parallel as part of the UK10K Project. Whereas there were no major differences in the allele frequency of common variants, a significant depletion of variants in the rare frequency spectrum was observed in Finns when comparing the two populations. On the other hand, we observed >2.1 million variants that were twice as frequent among Finns compared to Britons and 800,000 variants that were more than 10 times more frequent in Finns. Furthermore, in Finns we observed a relative proportional enrichment of variants in the minor allele frequency range between 2 - 5% (p < 2.2×10^−16^). When stratified by their functional annotations, loss-of-function (LoF) variants showed the highest proportional enrichment in Finns (p = 0.0291). In the noncoding part of the genome, variants in conserved regions (p = 0.002) and promoters (p = 0.01) were also significantly enriched in the Finnish samples. These functional categories represent the highest *a priori* power for downstream association studies of rare variants using population isolates.

## Introduction

Population isolates have not only provided insights into population diversity and history, but are also an exciting opportunity to identify rare and low-frequency variants associated with complex diseases^1–4^. So far, these studies have mostly focused on genetic variation in the coding regions with the highest enrichment observed in the variation that predictably disrupt protein coding genes.

Within coding regions, variant alleles that have high penetrance whilst predisposing to disease are likely to be deleterious and therefore kept at low frequencies by purifying selection in larger outbred populations^5–7^. Isolated populations resulting from recent bottlenecks have a substantial reduction in rare neutral variation and also many functional and even deleterious variants present at relatively higher frequencies because of increased drift and reduced selective pressure. Hence, recent isolates can be used to study causal variants that are rare in other populations in association with complex diseases^1–4^.

Finland is a well-known example of an isolated population where multiple historical bottlenecks resulting from consecutive founder effects have shaped the gene pool of current-day Finns^8^. Previous studies suggest the latest historical migration into Finland about 4000 years ago^9^. Due to lack of evidence of major migratory movements, it has been suggested that there were small but significant migrating groups of people.

Settlements resulting from the latter migratory movements mainly occurred along the south-east cost of Finland. Further, due to geopolitical reasons there have been additional major migratory movements within Finland in the 16th century in the eastern and northern parts of Finland. These settlements, initially founded by a small number of people, have grown in size over time leading to secondary population bottlenecks. An extreme example of the latter is Kuusamo, a county in the northeast part of Finland^10,11^. Historical records show that in 1718, there were 165 houses consisting of 615 individuals belonging to 39 families. Rapid population growth leading to a present day population of >15,000 individuals further increased the allelic drift in this sub-isolate. Consequences of these historical events have led to reduced genetic variation and higher overall linkage disequilibrium levels in Finland as compared to the outbred populations^10,11^.

During the last 1000 years, the Finnish population size has grown more than two orders of magnitude - from around 50,000 individuals to more than 5 million individuals. Furthermore, the most rapid growth has happened during the last 10 generations (~250 years), with population size growing from 500,000 to 5.4 million individuals. Combined with the historical bottleneck effect, these events have caused a massive departure from population genetic equilibrium whilst ‘shifting’ the proportion and frequency of many initially rare variants.

Such deviations have led to an increase in the prevalence of some monogenic Mendelian disorders in Finland as compared to the other parts of the world and are referred to as the Finnish disease heritage^12^ (FDH). Pronounced effects of the bottleneck have also been observed for complex diseases and disorders. For instance, schizophrenia is prevalent almost three times in northeastern sub-isolates as compared to rest of Finland^13^. Similarly, protective effects of enriched variants have also been observed as exemplified by variants in the LPA gene that protect against risk of cardiovascular diseases^1^.

However, the dynamics and properties of this genetic 'enrichment' are poorly understood on the genome scale, particularly outside the protein coding regions. We set out to provide a more comprehensive view of this enrichment and the other bottleneck effects in Finland by comparing whole-genome sequencing data in Finnish and British samples. In this study, we show how the historical bottlenecks have affected the genetic landscape of Finns and the frequency profile of variants across the entire genome. Whole-genome sequencing data gave us a unique opportunity to determine the enrichment of variants across both coding as well as non-coding regions of the human genome.

## Methods

### Sample selection

We sequenced the whole genomes of 1463 Finns at low coverage (~4.6×). These samples belonged to the FINRISK^14^ and H2000 cohorts. The FINRISK study comprises samples of the working age population, to study the risk factors associated with chronic diseases across Finland and is carried out every 5 years. The H2000 is a population based national survey aimed at studying the prevalence and determinants of important health problems amongst the working-age and the aged population (http://www.terveys2000.fi/julkaisut/baseline.pdf). Amongst these, 856 individuals have low HDL and 691 individuals have been diagnosed with psychosis. Further, 371 individuals belong to a sub-isolate within Finland, the Kuusamo region. Due to known genetic differences, for the comparison between Britons and the Finns, we restricted the analyses only to those Finnish individuals that are not from Kuusamo. To study the effects of a bottleneck within a bottleneck, 371 samples from non-Kuusamo Finns were further used for comparison against 371 samples from Kuusamo. All study participants gave their written informed consent to the study of origin.

### Whole genome sequencing and variant discovery

Low read-depth whole genome sequencing was performed at the Wellcome Trust Sanger Institute (WTSI). Joint variant calling of the raw binary sequence alignment map (BAM files) along with the UK10K samples was performed as part of the Haplotype Reference Consortium (HRC) (manuscript under review). The genotypes were further refined by re-phasing using SHAPEIT3 algorithm (O’Connell et al. 2015, under review). As a part of the joint calling quality control, only those sites that have a minor allele count of at least 5 copies in the entire dataset (32611 samples) went through additional filtering. Hence we have restricted the analyses to those variants with minor allele count >=5. The BAM files have been submitted to the European Genome-phenome Archive (EGA). In order to minimize the batch effects, we performed these analyses on only those British samples from UK10K (1463 samples from 3781 samples) that were also sequenced at the WTSI. We have only included autosomal single nucleotide variants for these analyses.

To determine the quality of the data, we compared the Finnish whole genome sequencing data with microarray genotypes for 629 individuals. From this comparison, we estimate that for variant sites with minor allele frequency >5% we observe 99.1% of variants with a low genotyping error of <0.1%. For the variant sites with minor allele frequency <5%, we estimate to have discovered 90.5% of the variants at an error rate of 6.9%.

### Annotations

The various functional categories were obtained as follows:

a. Coding sequence, promoters, untranslated region annotations were obtained from UCSC Genome Browser^15^ using the Gencode v19 gene models^16^.
b. Coding variants were further stratified using the Variant Effect Predictor^17^ into loss-of-function variants, missense variants and synonymous variants. Polyphen^18^ predictions were used to classify missense damaging variants.
c. Dnase1 hypersensitivity sites (DHS) were obtained from^19^ Trynka et al. 2013. We merged the coordinates for all cell types into one category.
d. Conserved regions in mammals were obtained from^20^ Linblad-Toh et al. 2011. These were post processed by Ward & Kellis^21^ 2012.
e. FANTOM5 enhancer coordinates were obtained from Andersson R et al.^22^ 2014
f. Super-enhancers were obtained from Hnisz et al.^23^ 2013. The genomic coordinates were merged over all cell types.
g. Transcription factor binding sites were obtained from Encode project^24^.

### Enrichment Analysis

We calculated the enrichment for each category beyond the baseline enrichment observed (enrichment calculated using all variants in Finns and Britons), assuming the following model.

Consider a category of variants in which we have observed F variants in the first population (e.g. Finnish) and B variants in the second population (e.g. British) and let M = F+B. Let s be the proportion of variants from the first population, and u the ratio of the numbers of variants in the first population to that in the second population. According to the binomial distribution, our point estimate for s is ŝ = F/M and has variance approximately ŝ(1-ŝ)/M. It follows that a point estimate for u is û = F/B, and the variance of log(û) is 1/(Mŝ(1-ŝ)) by the Delta method. This allows us to estimate 95% confidence intervals for û.

Suppose that we are comparing two categories of variants, and have observed F_1_ and B_1_ variants in category 1 and F_2_ and B_2_ variants in category 2. To test whether u_1_ is different from u_2_, we compute log(û_1_)-log(û_2_). Under the null hypothesis of no difference, this statistic has mean 0 and variance approximately (1/M_1_ + 1/M_2_)/(ŝ(1-ŝ)), where ŝ = (F_1_+F_2_)/(M_1_+M_2_), which we use to derive a p-value. Note that the standard proportion test between s_1_ and s_2_ gives essentially the same p-value.

## Results

### Overall frequency distribution of genome wide level variation

As a part of the SISu project, we sequenced the genomes of 1463 Finnish samples at low read-depth (average 4.6×) sampled across Finland. We compared these profiles to a sample of 1463 British individuals sequenced at average depth 7× as a part of the UK10K Consortium^25^. To reduce potential batch effects, these datasets were jointly processed as part of the Haplotype Reference Consortium^26^. After stringent quality control steps, we compared the minor allele frequencies (MAFs) of 10,457,802 and 11,172,232 single nucleotide variants (SNVs) identified with minor allele count 5 or greater in 1463 Finns and in the same number of Britons, respectively (Table 1).

**Table 1:**
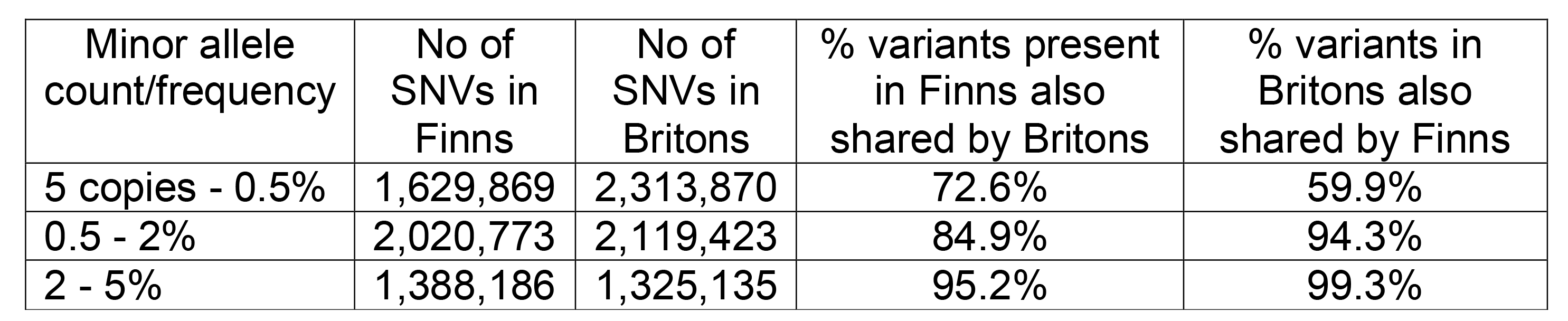
Summary of SNVs studied in Finnish and British samples.

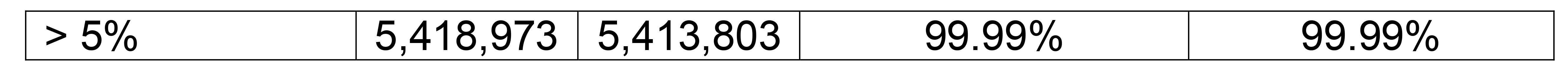

As a direct result of the bottleneck effect, we observed that Finns have significantly fewer rare variants (MAF < 0.5%) compared to Britons (Figure 1). On the other hand, in Finns, we determined proportionally small but significant enrichment of low-frequency variants (MAF range between 2 - 5%, binomial p < 2.2×10^−16^). The latter is also a direct effect of the historical bottleneck, followed by population growth. And as expected, we observed no differences in the number of common (MAF > 5%) variants (Figure 1).

**Figure 1:**
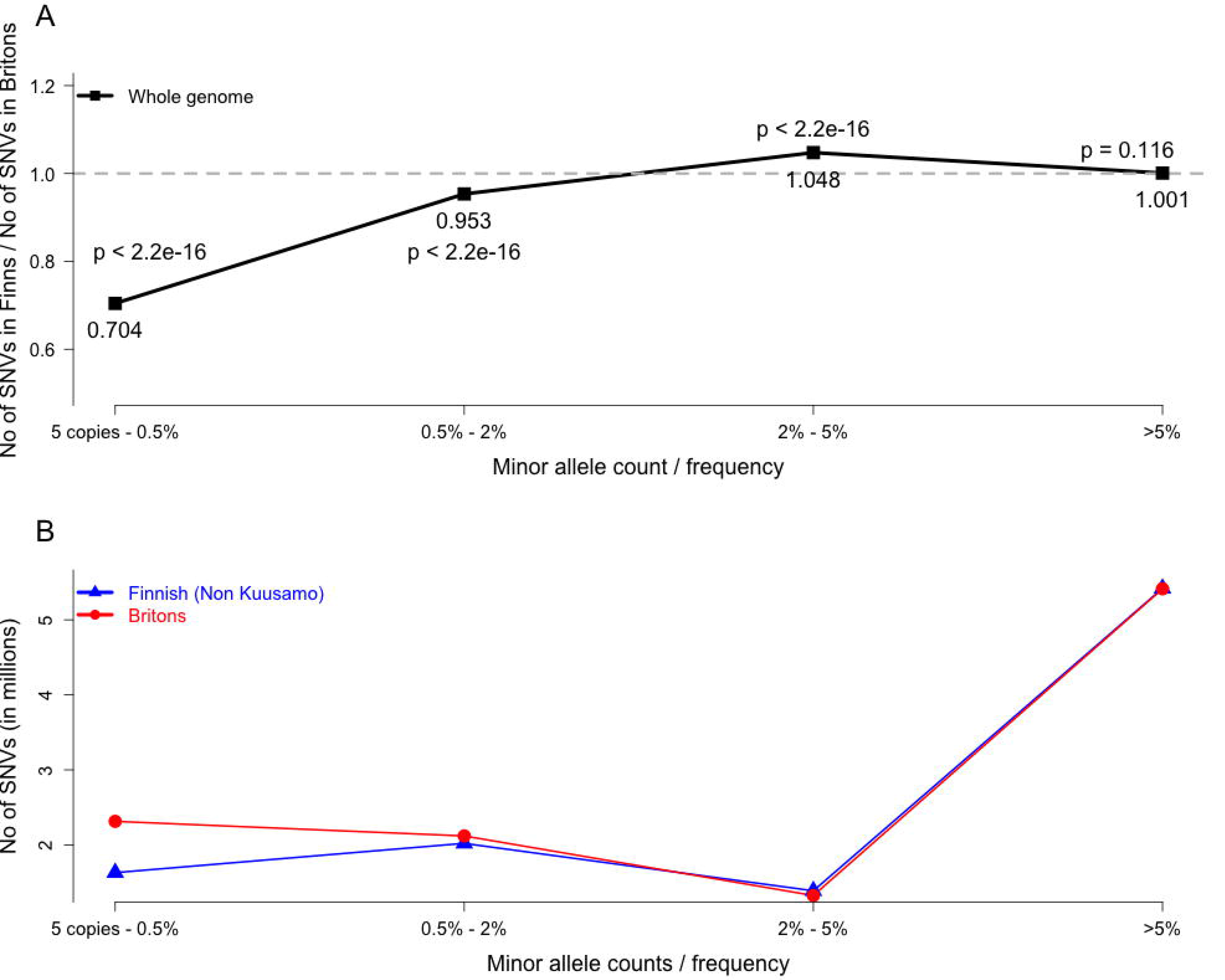
(A) Allele frequency spectrum of variants across the whole genome in Finns compared to the Britons. The black line represents the ratio of the number of variants observed in Finns to those in Britons. (B) The number of variants seen in each population across the genome in different MAF bins. The lines in blue and red represent the number of variants for each bin observed in Finns and Britons, respectively.

For each frequency range, we also calculated the percentage of variants shared between both population samples. As anticipated, the number of variants observed as rare in Britons (MAF < 0.5%) and also found as polymorphic in Finns was considerably lower than the opposite: only 54.7% of variants with MAF < 0.5% in Britons were polymorphic in Finns while 72% of variants with MAF < 0.5% in Finns were also polymorphic in Britons (Figure 2). However, for the MAF range of 0.5 - 5% the opposite was true: a lower proportion of variants seen in Finns were also polymorphic in Britons (e.g. for 0.5 - 2% range, 84.9% and 94.3% of variants are shared, respectively). For common variants (MAF > 5%), essentially all (99.9%) were observed to be shared in both directions (Table 1 and Figure 2).

**Figure 2:**
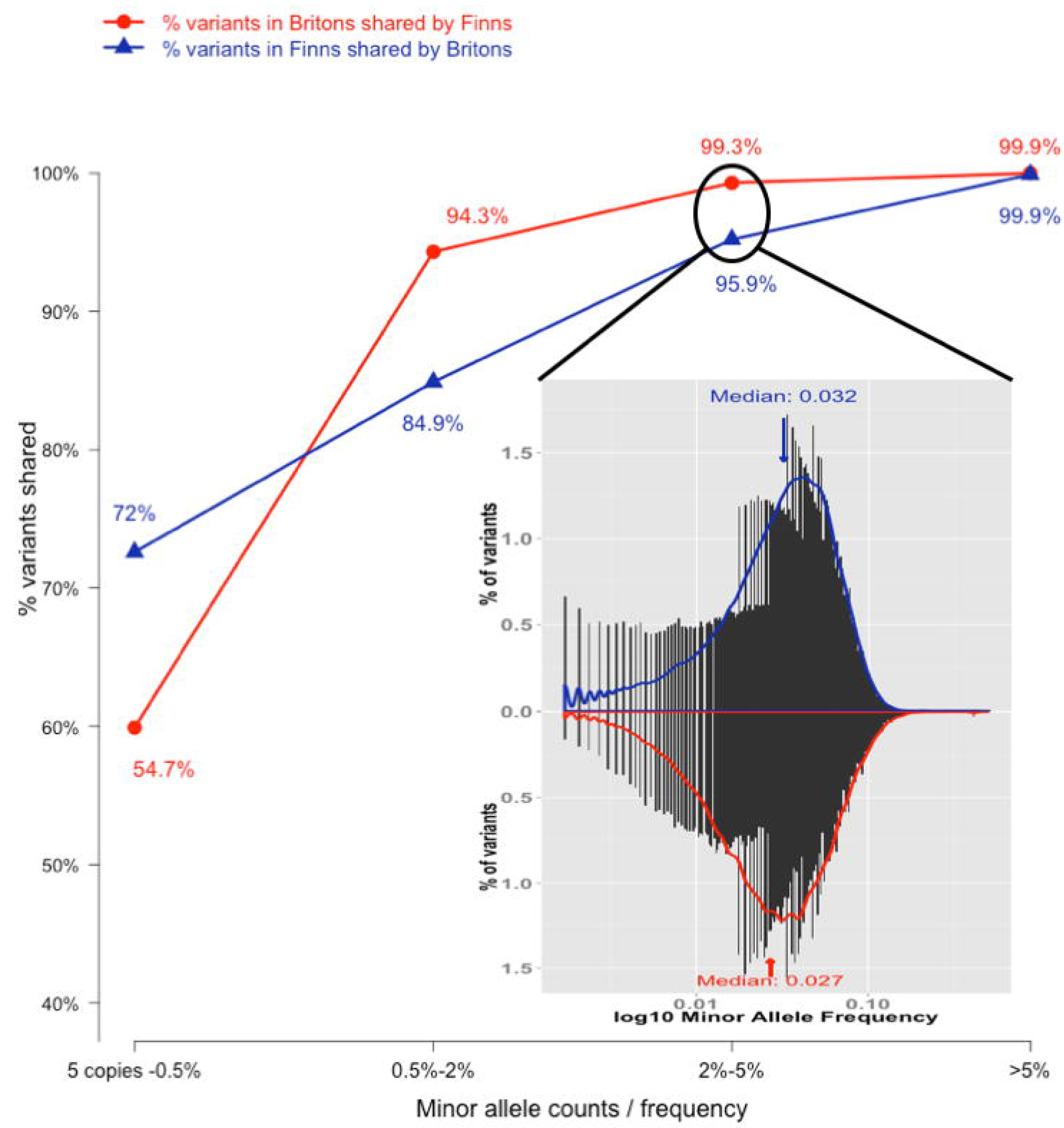
Variants shared between the two populations. The percentage of variants that are shared between the Finns and the Britons across different allele frequency bins. The histograms represent the allele frequencies of the shared variants in the other population for the MAF bin 2 - 5%.

### Enrichment of variants across functional categories

We also calculated the relative enrichment of Finnish SNVs across various functional categories shown to be relevant in different phenotypic traits including disease^27^. For each of these categories, we compared its distribution profile with that of the ‘expected’ whole genome baseline distribution in Finns (Figure 1A). Although there were several small deviations from the expected baseline in almost all functional categories, the greatest differences were consistently observed in the MAF range of 2 - 5% (Supplementary Figures 1-8).

In accordance with the latter observation, we compared the enrichment of different functional categories for MAF range 2 - 5% (Figure 3). Across studied functional categories, the coding regions showed the highest enrichment in Finns (Figure 3). More specifically, we observed > 1.3-fold enrichment of loss-of-function (p = 0.0291) variants and > 1.1-fold enrichment of missense (p = 0.0197) variants (Figure 4A), similarly as was demonstrated previously in Finns by exome sequencing^1^. Furthermore, we observed consistent enrichment of rare and low-frequency (MAF ≤ 5%) missense damaging variants (Figure 4A).

**Figure 3:**
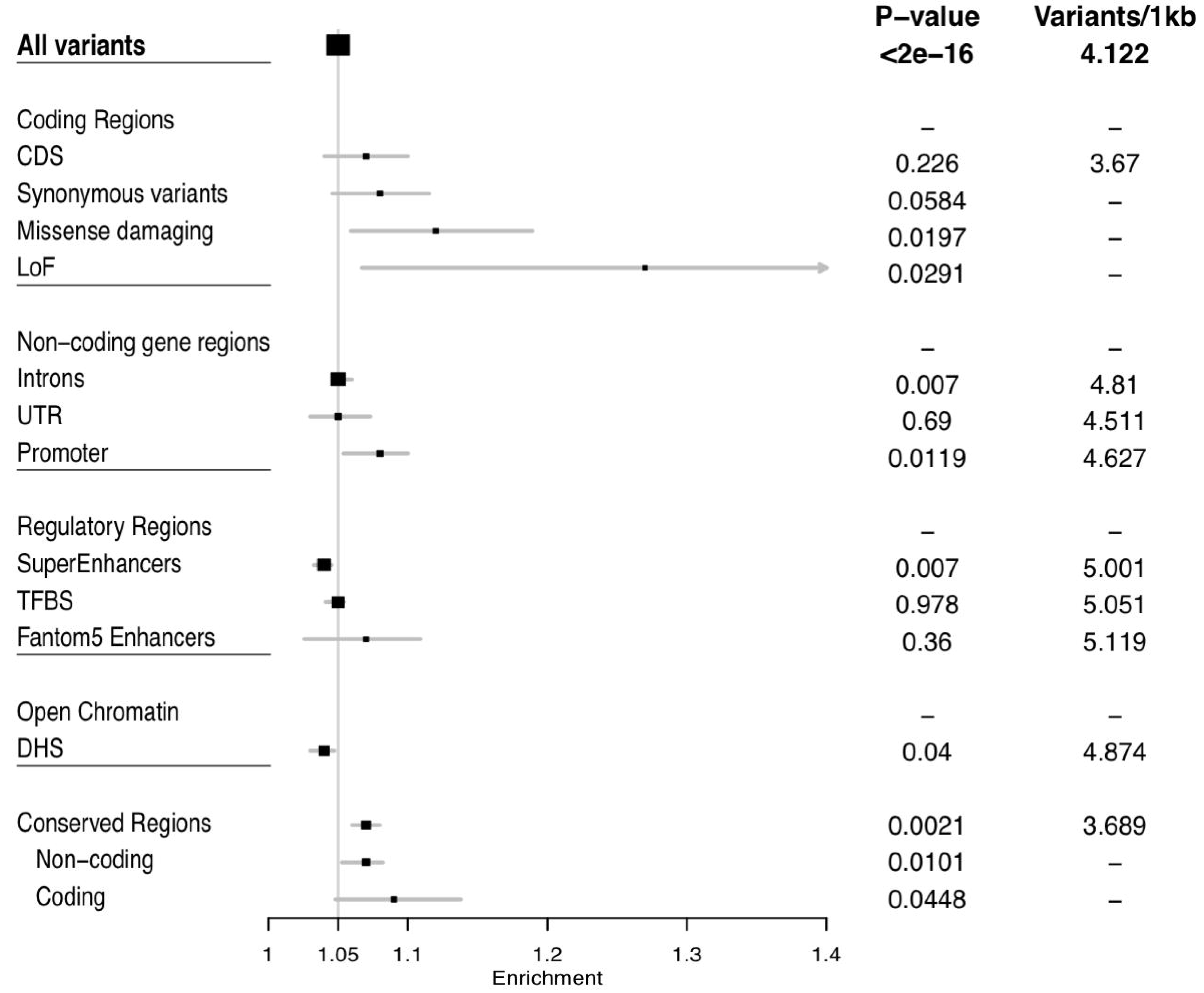
Forest plot showing the enrichment across various functional categories for the variants in the minor allele frequency range 2 - 5%, where we observe consistent enrichment across most categories). The sizes of the boxes correspond to the size of each category and the black horizontal lines represent the 95% confidence intervals. Proportional enrichment is calculated compared to Britons.

**Figure 4:**
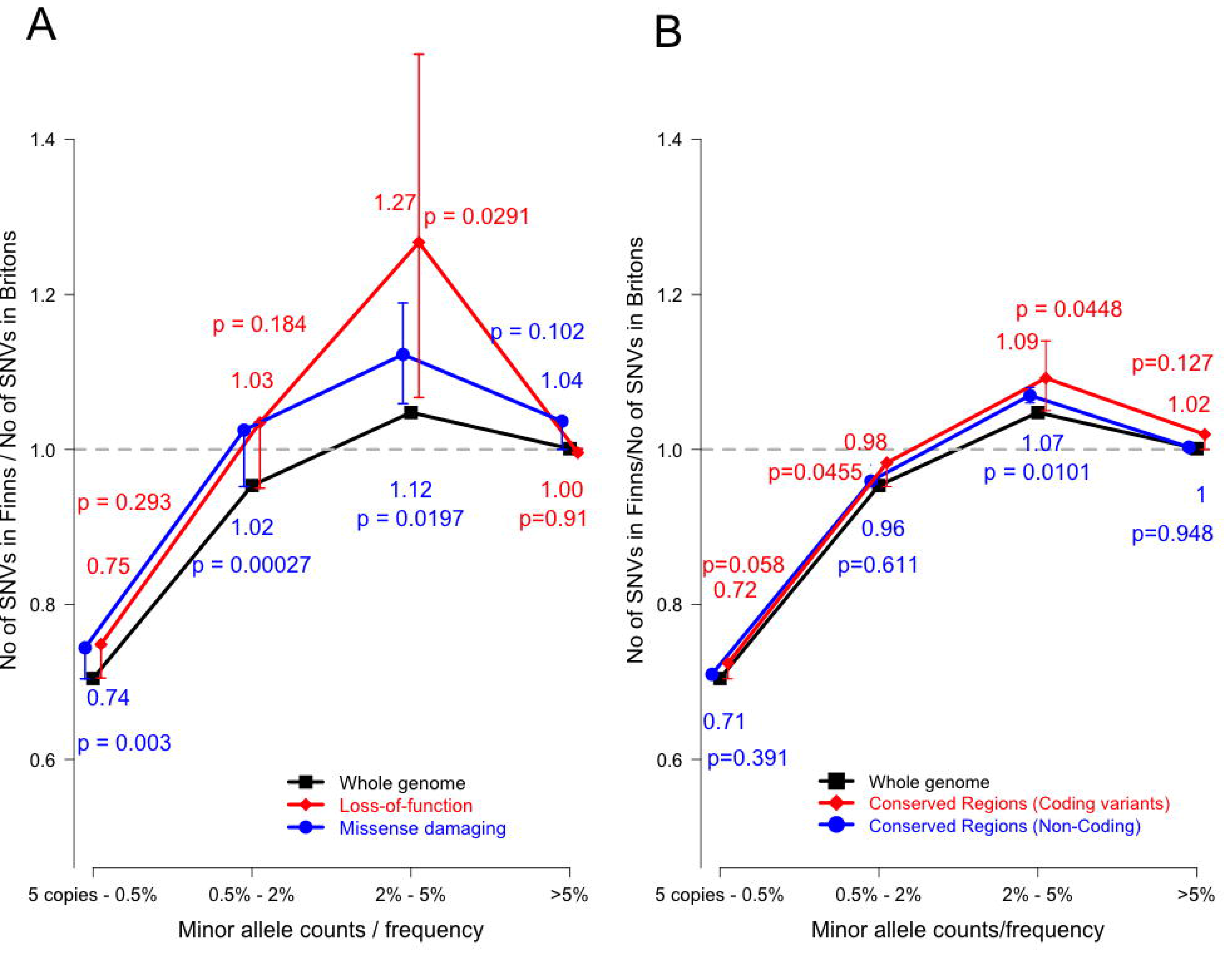
Comparison of variants across different allele frequency spectrums in functional categories. (A) Proportional enrichment of LoFs in Finns compared to Britons. The red line represents the ratio of the number of LoF variants in Finns compared to Britons. The black line shows the baseline enrichment observed across the whole genome. (B) Proportional enrichment of the number of variants in the conserved regions in Finns compared to Britons. The red line represents the variants common between conserved regions and the coding regions. The blue line represents the variants in the conserved regions but not in the coding regions. The black line shows the baseline enrichment observed across the whole genome.

As observed for the low-frequency variants (MAF 2 - 5%) in the coding regions, we found enrichment in the non-coding regions as well. In the non-coding parts of the genome, the promoter regions showed the largest enrichment compared to the expected baseline (p = 0.012, Supplementary Figure 3), followed by the conserved non-coding regions of the human genome (p = 0.01, Figure 4B). Although the Fantom5 enhancer regions showed proportional enrichment, it was not significant compared to the expected baseline (Figure 3; Supplementary Figure 4). The other functional categories followed the baseline enrichment (Figure 3). We also observed that, although enriched when compared to the Britons, the DNase1 Hypersensitivity sites (DHS) and the super-enhancer elements are only marginally depleted beyond the expected bottleneck effects (p_DHS_ = 0.04 and p_super-enhancers_ = 0.007, Supplementary Figures 5 and 7).

### MAF-enrichment of variants and effect on statistical power

We observed that 20.16% of all variants in Finns have minor allele frequencies elevated at least two-fold. Furthermore, 1.36% of these variants were enriched >=50 fold. For the proportionally enriched functional categories, we calculated the number of variants with elevated frequencies in Finns as compared to Britons (Table 2) and observed even higher MAF-enrichment for many of these categories. Missense damaging variants showed the highest enrichment with 37.98% variants showing minor allele frequencies at least twice as high as observed in the Britons. 29.71% of the loss-of-function variants showed at least two-fold MAF-enrichment compared to the British sample.

**Table 2:** MAF-enrichment of variants in Finns and Britons. This table summarizes the number of variants that are MAF-enriched in both populations across the whole genome and the categories in which there is a significant enrichment. The first column describes the fold-change of minor allele frequency between the two samples.

| Finns |  |  |  |  |  |  |
| --- | --- | --- | --- | --- | --- | --- |
| Enrichment | Genome-wide | LoF | Missense-damaging | Conserved Regions-Coding | Conserved Regions-Non Coding | Promoters |
| 2-5x | 722587 (6.91%) | 177 (7.815%) | 1704 (9.936%) | 3109 (8.681%) | 22823 (7.665%) | 9179 (7.196%) |
| 5-10x | 561243 (5.367%) | 225 (9.934%) | 2083 (12.146%) | 3013 (8.413%) | 19385 (6.51%) | 7415 (5.813%) |
| 10-50x | 682275 (6.524%) | 233 (10.287%) | 2360 (13.761%) | 3643 (10.173%) | 23207 (7.794%) | 9129 (7.157%) |
| >=50x | 142354 (1.361%) | 38 (1.678%) | 366 (2.134%) | 655 (1.829%) | 4584 (1.539%) | 1888 (1.48%) |
| Britons |  |  |  |  |  |  |
| Enrichment | Genome-wide | LoF | Missense-damaging | Conserved Regions-Coding | Conserved Regions-Non Coding | Promoters |
| 2-5x | 935392 (8.383%) | 203 (8.547%) | 1839 (9.98%) | 3483 (9.07%) | 28862 (8.987%) | 11242 (8.21%) |
| 5-10x | 1319454 (11.824%) | 405 (17.053%) | 3991 (21.658%) | 6336 (16.5%) | 44903 (13.982%) | 17498 (12.778%) |
| 10-50x | 893522 (8.007%) | 235 (9.895%) | 2447 (13.279%) | 4005 (10.429%) | 29368 (9.145%) | 11557 (8.44%) |
| >=50x | 25552 (0.229%) | 12 (0.505%) | 58 (0.315%) | 125 (0.326%) | 818 (0.255%) | 382 (0.279%) |

We also performed the same analyses in Britons. Across all categories, for variants that are enriched at the most 10 fold, the Britons consistently show much higher number of variants across all enriched functional categories. However, for the loss-of-function and missense damaging categories beyond 10-fold enrichment, the Finns have larger proportion of variants enriched. Interestingly, beyond 50-fold enrichment, the Finns have a relatively larger proportion of variants enriched across all functional categories (Table 2).

This enrichment of minor allele frequencies for certain variants boosts the statistical power to detect possible associations with traits and diseases. To quantify this gain in power, we calculated the number of samples required to detect association with high probability in Finns as compared to the Britons (Figure 5). For variants that are twice as common among Finns to Britons, only half the number of individuals would be required to detect the associations in Finnish samples. Further, for variants with minor allele frequencies enriched 5× and 10× times, only 20% and 10% of the samples respectively are required to detect associations. These analyses indicate the gain in power for studying association analyses in isolated populations such as the Finnish population.

**Figure 5:**
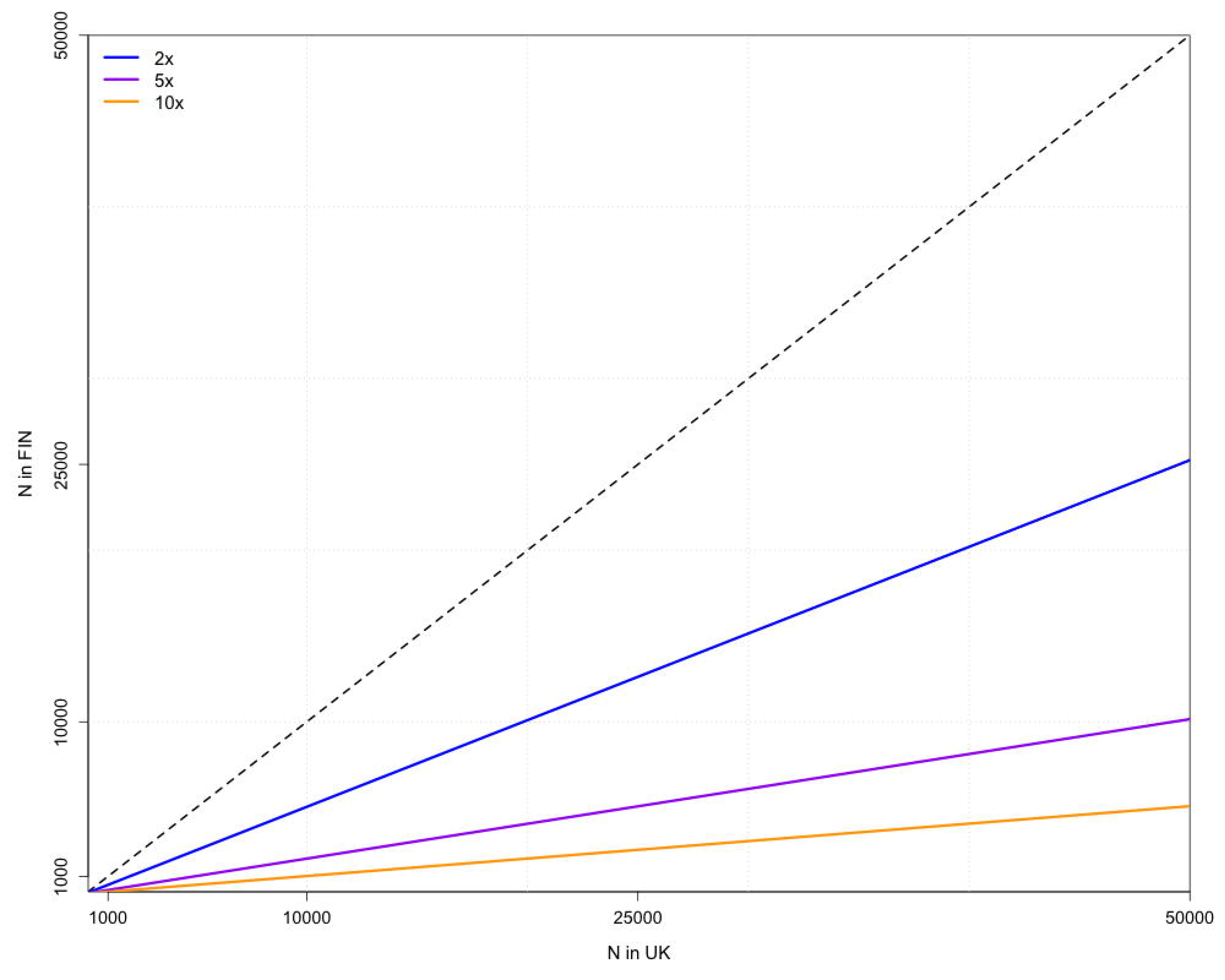
Statistical power gained due to enrichment of a variant in Finns. Plot showing the number of Finnish samples required to detect association if a variant is enriched two fold (blue), five-fold (purple) and ten-fold (yellow) in Finns as compared to Britons.

### Sub-isolate of an isolated population

Amongst the sequenced Finnish individuals, 371 belonged to the Kuusamo sub-isolate within Finland. When comparing the SNV frequency profiles of these individuals against the same number of randomly selected non-Kuusamo Finns, we observed a significant reduction in the number of rare variants (Supplementary Figure 9). Although there was no overall enrichment of low-frequency variants, when looking at the variants stratified by their functional categories, we found a significant enrichment of LoF variants in the MAF 0.5 - 2% frequency range (p = 0.0272) (Supplementary Figure 10).

## Discussion

Studying relatively recently bottlenecked and isolated populations or sub-isolates provides an excellent opportunity to discover disease-associated genes, as some of the underlying (and initially rare) variants can reach much higher frequency after the population bottleneck. We studied this bottleneck effect and subsequent enrichment of variants in Finnish samples by comparing them to outbred British samples. We demonstrated how the historical bottlenecks have affected the genetic landscape of Finns and the frequency profile of variants across the entire genome.

As expected, we observed no major differences in the common variant frequency spectrum – as most variants with MAF > 5% probably segregated already tens of thousands of years ago, they are known to be relatively equally distributed in populations that separated more recently^28,29^. On the other hand, there was a significant depletion of variants in the rare frequency spectrum in Finns. Also, as an additional hallmark of the population bottleneck, a significant enrichment of low-frequency variants was observed (Figure 1). For most functional variants we observe an enrichment beyond the expected baseline showing that bottleneck population have a higher likelihood of accumulating deleterious and disease associated mutations. In order to test the robustness of enrichment of low frequency variants, we changed the minor allele frequency bins for the whole genome analysis. We observed consistency in the enrichment of low frequency variants (MAF 1 - 5%; Supplementary Figure 11). This phenomenon also explains the high prevalence of several monogenic Mendelian disorders, so-called ‘Finnish disease heritage’, caused by genetic disease variants found at much higher frequencies in Finland than in the rest of the Europe^12^.

We observed that within the frequency range of MAF 0.5 - 2%, only a subset (84.9%) of the variants in Finnish samples is also seen in the British samples (Figure 2). For the common variants, in contrast, most variants (99.9%) were shared between the two populations (Figure 2). These findings are similar to the patterns observed in the Icelandic^3^ and the Sardinian populations^2^. Finns also show a similar enrichment of LoF variants and missense variants as seen in the Icelandic populations. However, the enrichment observed in the Icelandic population was found in the lower minor allele frequency range as opposed to the Finnish sample, possibly due to the differences in the historical bottleneck ‘width’, time since bottleneck (the Icelandic bottleneck was more recent than the Finnish), and the subsequent population growth rate. As such, this enrichment can provide a boost in statistical power when studying health-related phenotype traits affected by these enriched variants.

Other studies have recently demonstrated that functional categories such as conserved regions and Fantom5 enhancers contribute disproportionately more to the heritability of complex diseases, suggesting that in addition to coding regions also regulatory regions are enriched for trait and disease associated variation^27,30,31^. Here, we used 12 functional annotations to determine if variants in any of these categories are enriched beyond the baseline distribution of variants (and bottleneck effect) in Finns. We observed an enrichment across most functional categories in the low-frequency bin (MAF 2 - 5%). As reported previously^1^, we observed a significant enrichment of low-frequency LoF and missense variants in Finns (Figure 4A). In addition to the enrichment of coding variants, however, also non-coding conserved regions and non-coding genic regions such as intron and promoter regions showed enrichment beyond the baseline bottleneck effect (Figure 4B). This enrichment likely appears due to selection against these variants in noncoding conserved regions and non-coding genic regions in outbred European populations. Furthermore, we see depletion for the super-enhancer regions and the DHS elements. This suggests that functionally, super-enhancers may be actually less active than regular enhancers, as was also proposed previously ^27,32^.

Previous studies have shown the utility of bottleneck populations to identify variants with elevated frequencies to identify associations with diseases and phenotypes^1,2,13^. Our findings show that across the genome, around 20% of all variants present in Finns have enrichment at least twice as observed in the Britons (Table 2). The percentage of variants with at least 2× enrichment further increases for loss-of-function variants and missense damaging variants (29.71% and 37.98% respectively). Our power calculation simulations show that by testing for associations with these variants, the number of samples required to achieve significant detections are much lower (Figure 5). These analyses suggest the great value of testing for genetic associations due to the gain in statistical power in isolated populations such as Finns.

Although we tried to eliminate all possible sources of biases and other technical limitations by jointly processing our datasets, our results might be somewhat limited in the very rare variant spectrum. FINRISK and Health2000 cohorts have collected samples from all over mainland Finland. In this study, however, the samples have been geographically randomly selected. As low-coverage whole-genome sequencing data are sub-optimal for detecting variants observed only in a few individuals, rare variants observed in Britons were likely to be called more confidently compared to similar variants in Finns. Additionally, the British dataset had slightly higher coverage than the Finnish data (4.6× vs. 7×), which may have had some effect on calling of the rare and low-frequency variants in Finns. Such technical limitations and differences may have led to under-estimation of our main findings (except the depletion of rare variants in Finns). Our comparison was limited against British samples and autosomal SNVs only, and future studies should therefore carry out comparisons against a panel of jointly processed heterogeneous population samples, including all types of variants (also from sex chromosomes). When comparing the Kuusamo sub-isolate sample to the Finnish non-Kuusamo individuals, we found that only LoF variants (that also showed the largest enrichment between Finns and Britons) appear significantly enriched. This is possibly due to the small sample size of the Kuusamo subset.

This study provides insights into the effects of a population bottleneck in various functional categories across the whole human genome. Obvious advantages of isolated populations are significantly reduced heterogeneity in genetic architecture, phenotype and the environment. The frequency of an originally rare allele that passed through the population bottleneck can be increased by several orders of magnitude (even >100-fold for some variants), after which it will decline relatively slowly (due to selective pressure). This phenomenon will therefore increase the statistical power to identify rare variants associated with complex disorders in both coding as well as noncoding regions of the human genome in isolated populations^1,13^.

## Acknowledgements

This work was supported by the Doctoral Programme in Biomedicine-DPBM (H.C.), the Academy of Finland (251217, 255847 and 285380 to S.R.; 251704 and 286500 to A.P.; 295504, 269862 to T.A.; 257654 and 288509 to M.P.), the Academy of Finland Center of Excellence for Complex Disease Genetics of the Academy of Finland (grant nos. 213506 and 129680), the Wellcome Trust (WT089062/Z/09/Z and WT089061/Z/09/Z to S.R., and WT098051 for S.M, K.W. and R.D.), EU FP7 projects ENGAGE (201413 to A.P. & S.R.), BioSHaRE (261433 to A.P. and S.R.), the Finnish Foundation for Cardiovascular Research (A.P., S.R. and V.S.); the Sigrid Juselius Foundation, Biocentrum Helsinki (S.R.), the Nordic Information for Action eScience Center (NIASC)-a Nordic Center of Excellence financed by NordForsk (grant no. 62721 to A.P., P.P. & S.R.); IUT20-60 Omics for health: an integrated approach to understand and predict human disease (P.P). The authors also wish to acknowledge CSC - IT Centre for Science, Finland and the FIMM Technology Centre services for computational resources.

